# Cell-autonomous regulation of structural and functional plasticity in inhibitory neurons by excitatory synaptic inputs

**DOI:** 10.1101/240176

**Authors:** Hai-yan He, Wanhua Shen, Lijun Zheng, Xia Guo, Hollis T. Cline

## Abstract

Functional circuit assembly is thought to require coordinated development of excitation and inhibition, but whether they are co-regulated cell-autonomously remains unclear. We investigated effects of decreased glutamatergic synaptic input on inhibitory synapses by expressing AMPAR subunit, GluA1 and GluA2, C-terminal peptides (GluA1CTP and GluA2CTP) in developing Xenopus tectal neurons. GluACTP decreased excitatory synaptic inputs and cell-autonomously decreased inhibitory synaptic inputs in excitatory and inhibitory neurons. Visually-evoked excitatory and inhibitory currents decreased proportionately, maintaining excitation/inhibition. GluACTP affected dendrite structure and visual experience-dependent structural plasticity differently in excitatory and inhibitory neurons. Deficits in excitatory and inhibitory synaptic transmission and experience-dependent plasticity manifested in altered visual receptive field properties. Both visual avoidance behavior and learning-induced behavioral plasticity were impaired, suggesting that maintaining excitation/inhibition alone is insufficient to preserve circuit function. We demonstrate that excitatory synaptic dysfunction in individual neurons cell-autonomously decreases inhibitory inputs and disrupts neuronal and circuit plasticity, information processing and learning.

---

Activity plays a critical role in the refinement and maintenance of functional neural circuits^1^, which is thought to require coordinated development of its two principle components: excitatory and inhibitory neurons^2^. Although proportional co-regulation of excitation and inhibition and a constant excitation/inhibition ratio have been widely observed during circuit development^2^ our understanding of how glutamatergic excitatory inputs affect the development of inhibition at synaptic and neuronal levels remains incomplete. Mounting evidence from different brain regions and species suggests that perturbing activity or sensory experience decreases development of inhibition and disrupts the maturation and specification of inhibitory neurons and circuits^3–7^, however most of these studies perturbed activity broadly and were unable to resolve cell-autonomous and circuit-based outcomes. Direct evidence that glutamatergic synaptic inputs drive the cell autonomous development of inhibitory input in individual neurons is unclear.

As the predominant mediator of fast excitatory synaptic transmission, AMPARs provide the initial depolarization that is essential for the activation of NMDARs and subsequent secondary signal transduction and synaptic plasticity mechanisms. Four types of AMPAR subunit (GluA1–4) form different hetero- and homo-dimers of AMARs, with GluA1 and GluA2 being the major AMPAR subunits. Regulation of the trafficking of postsynaptic AMPAR underlies activity-dependent plasticity of synaptic strength^8–10^. Regulatory sites within the C-terminal region of GluA1 and GluA2 subunits are required for synaptic trafficking of AMPARs^8,11^. Expression of peptides corresponding to the GluA C-terminal peptides (GluA1CTP or GluA2CTP) impairs AMPAR trafficking, decreases excitatory synaptic transmission and disrupts experience-dependent synaptic plasticity^12–14^ GluACTPs are therefore effective tools to disrupt AMPAR-mediated excitatory transmission in individual neurons, permitting study of outstanding questions concerning the role of excitatory synaptic inputs in structural and functional development of neurons and circuits.

Here, we expressed GluA1CTP or GluA2CTP, referred to collectively as GluACTPs, in individual tectal neurons to assess the effects of impaired excitatory synaptic transmission on inhibitory synaptic inputs and the development of structural and functional properties in excitatory and inhibitory neurons in vivo. We show that GluACTP expression proportionally decreased excitatory and inhibitory synaptic inputs, resulting in a constant balance of excitation to inhibition in both inhibitory and excitatory neurons. In vivo time-lapse imaging demonstrated that deficits in excitatory synaptic inputs had distinct effects on dendritic arbor development and experience-dependent structural plasticity in excitatory and inhibitory neurons. GluACTP-mediated decreases in excitatory and inhibitory transmission also manifested in deficits in visual information processing, recorded as impaired spatial and temporal receptive field properties, and visuomotor behavior. Finally, GluACTP expression blocked learning-induced behavioral plasticity. Our results demonstrated that excitatory synaptic dysfunction led to cell-autonomous inhibitory synaptic dysfunction, which then ramified to impair neuronal and circuit properties and degrade behavioral performance.

## Results

### GluACTP expression reduced both excitatory and inhibitory synaptic transmission in tectal neurons

To test whether decreasing glutamatergic synaptic inputs in individual neurons affects GABAergic synaptic transmission, we sparsely transfected tectal neurons with constructs co-expressing GFP and GluA1CTP or GluA2CTP and recorded mEPSCs and mIPSCs from GFP+ neurons 5–8 days later (Fig. 1a). mEPSC frequency was significantly reduced in both GluA1CTP and GluA2CTP expressing neurons, with no significant change in mEPSC amplitudes (Fig. 1b-c). The decrease in mEPSC frequency likely reflects loss of synapses over several days of GluACTP expression. Interestingly, both the frequency and amplitude of mIPSCs were significantly reduced in GluACTP-expressing neurons (Fig. 1d-e), suggesting that excitatory synaptic inputs govern the development of inhibitory synaptic inputs in a cell-autonomous manner. By contrast, disrupting inhibitory synaptic inputs by interfering with GABA_A_R trafficking does not affect excitatory input onto the same neurons^15^. Paired pulse ratios of excitatory synaptic currents were comparable in neurons expressing GluA1CTP (n=5), GluA2CTP (n=5) or controls (n=7) (Fig. 1f-g), consistent with a deficit in AMPAR trafficking into postsynaptic sites^12–14,16^.

**Figure 1.**
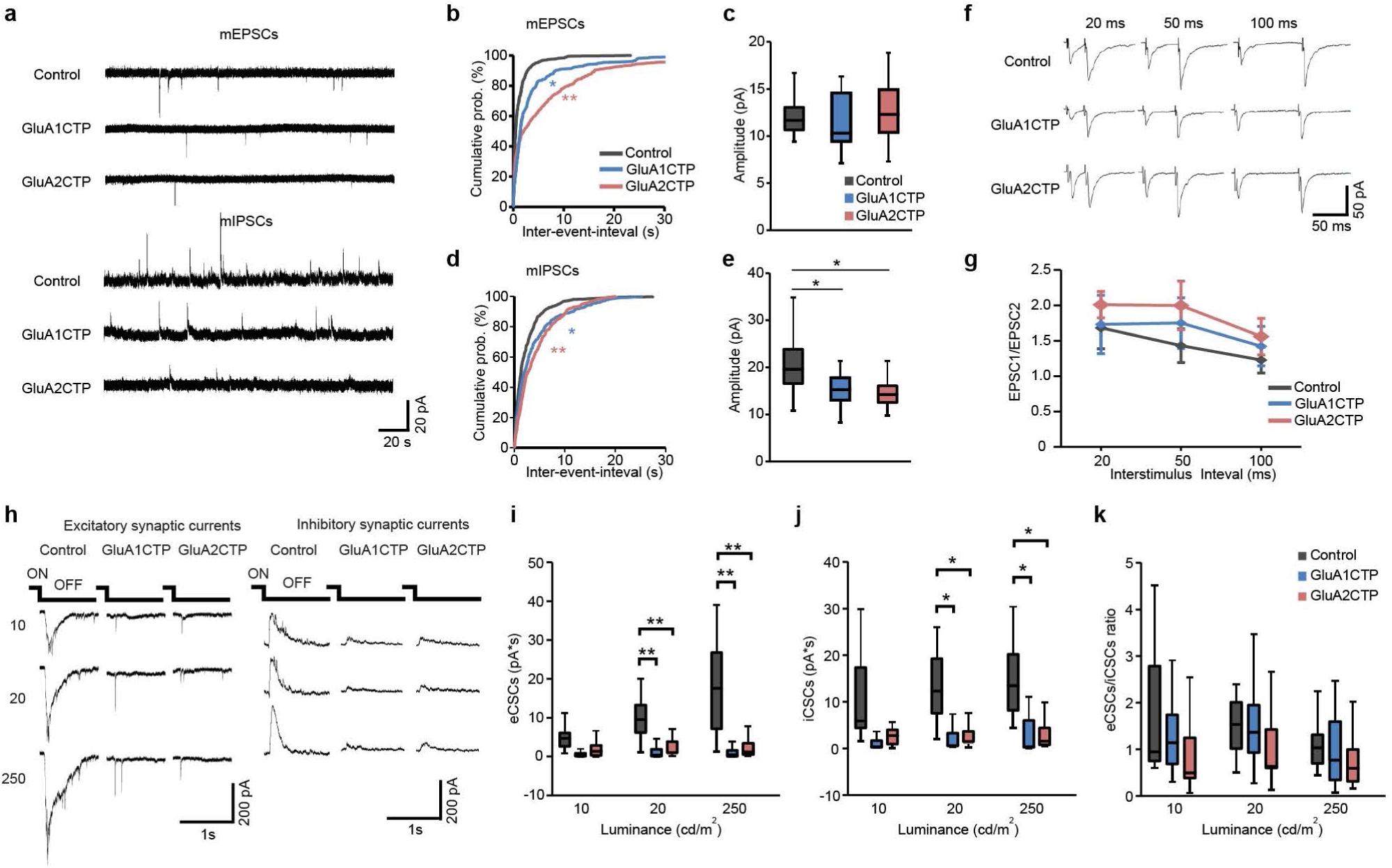
GluACTP expression reduced both spontaneous and evoked excitatory and inhibitory synaptic transmission in tectal neurons. (**a**) Representative traces of mEPSCs and mIPSCs for neurons expressing EGFP only (Control), GluA1CTP and GluA2CTP respectively. (**b**) Expression of GluA1CTP or GluA2CTP significantly increased interevent intervals (IEIs) of mEPSCs in tectal neurons. * *P* <0.05, ** *P* < 0.01. (**c**) Amplitudes of mEPSCs were not significantly affected by GluA1CTP or GluA2CTP expression. Control: n=14; GluA1CTP: n=13; GluA2CTP: n=15. (**d-e**) Cumulative distributions (**d**) and amplitudes (***e***) of mIPSCs showing GluA1CTP or GluA2CTP expression significantly increased IEI and decreased amplitudes of mIPSCs compared to control neurons. \**P*<0.05, **P<0.01. (**f**) Representative recordings of EPSCs in response to paired stimuli 20, 50 and 100 ms apart in neurons from each experimental group. Stimulus artifact was clipped for clarity. (**g**) Pair-pulse ratios of EPSC2/EPSC1 were not significantly different between control, GluA1CTP- and GluA2CTP-expressing neurons. Scale bar: 20 pA, 20 ms. Control: n=7; GluA1CTP: n=5; GluA2CTP: n=5. (**h**) Representative traces for visually-evoked excitatory CSCs (eCSCs) and inhibitory CSCs (iCSCs) in control, GluA1CTP- and GluA2CTP-expressing neurons in response to full field light off visual stimuli at intensities of 10, 20 and 250 CD/cm^2^. (**i-j**) Summary data showing that eCSCs (**i**) and iCSCs (**j**) in GluA1CTP or GluA2CTP-expressing neurons are significantly decreased compared to control neurons in response to visual stimuli of 20 and 250 CD/cm^2^ respectively. Control: n=7; GluA1CTP: n=7; GluA2CTP: n=7. (**k**) The ratio of iCSCs to eCSCs in GluA1CTP- and GluA2CTP-expressing neurons remained comparable to control neurons in response to visual stimulation of all luminances tested.

Tectal neurons receive direct excitatory retinal inputs as well as feed forward and feedback inhibitory inputs within tectal circuits^10,17,18^. To determine if the decreased excitatory synaptic transmission affected the E/I balance of evoked synaptic responses, we recorded excitatory and inhibitory compound synaptic currents (eCSCs and iCSCs) from transfected tectal neurons evoked by full field visual stimulation in intact animals (Fig. 1h). Visually-evoked eCSCs and iCSCs recorded from either GluA1CTP or GluA2CTP-expressing neurons were significantly smaller than controls (Fig. 1i-j), however, the ratio of total integrated charge transfer between iCSCs and eCSCs remained unchanged between GluACTP-expressing and control neurons (Fig. 1k). These data further demonstrate that interfering with GluA1- and GluA2-containing AMPAR trafficking not only decreases excitatory synaptic transmission onto the transfected cells, but also induces a proportional decrease in the inhibitory synaptic transmission onto the same neurons.

### Decreased excitatory inputs induced cell-autonomous decreases in inhibitory synaptic inputs in both excitatory and inhibitory neurons

Excitatory and inhibitory tectal neurons demonstrate different visual experience-dependent structural and functional plasticity^19^. To test whether disrupting excitatory synaptic transmission affects excitatory and inhibitory neurons differentially, we combined immunohistochemical labeling of the excitatory and inhibitory synaptic markers PSD95 and gephyrin, with GABA immunolabeling to examine the density of excitatory and inhibitory postsynaptic puncta in dendrites of GluACTP-expressing neurons.

In the optic tectum, individual tectal neurons express a mixture of AMPARs with different subunit compositions (GluA2-lacking or GluA2-containing). The proportion of GluA2-lacking to GluA2-containing AMPARs varies among individual tectal neurons such that more immature neurons show higher content of calcium permeable GluA2-lacking-AMPARs^20^. Both excitatory and inhibitory neurons express GluA1 and GluA2 in the developing optic tectum, as shown by double immunolabeling with GABA and GluA1 or GluA2 antibodies (Fig.2a-b).

**Figure 2.**
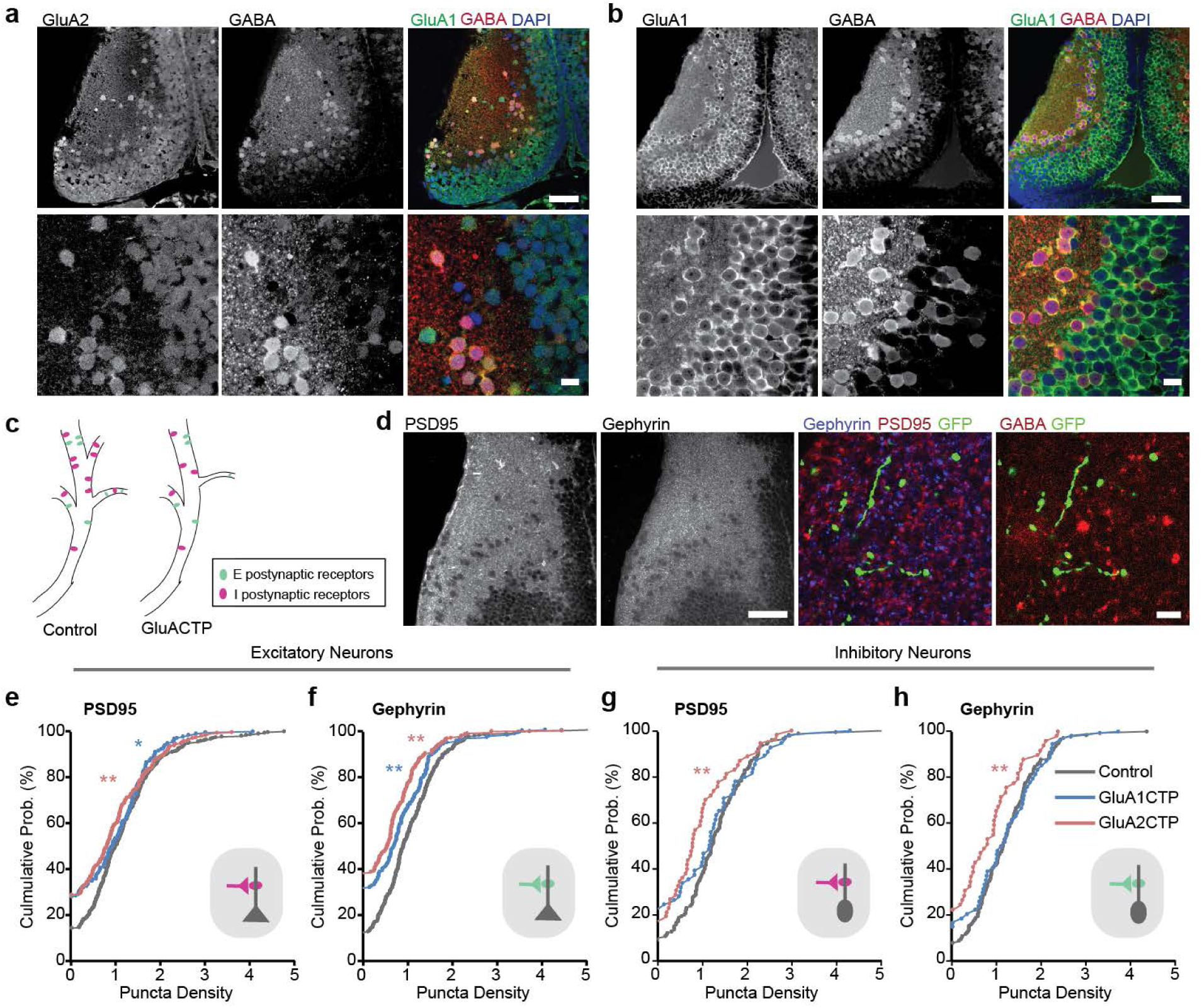
Decreased excitatory inputs induced cell-autonomous decreases in inhibitory synaptic inputs in both excitatory and inhibitory neurons. (**a-b**) Co-immuno labeling of GABA and GluA2 (**a**) or GluA1 (**b**) antibodies shows that both GluA1 and GluA2 are widely expressed in the tectum and are found in both excitatory (GABA-negative) and inhibitory (GABA-positive) neurons. Scale bar: top: 50μm; bottom: 10μm. (**c**) Schematic illustrates excitatory and inhibitory synapses on excitatory and inhibitory postsynaptic neurons. (**d**) Psd95 and gephyrin immunolabeling in the tectum (left) show high puncta density in the neuropil and relatively low density in the somatic region. Right: representative images of PSD95 and gephyrin puncta in a GFP+ dendritic segment. The identity (excitatory or inhibitory) of the GFP+ dendrite was determined by GABA immunolabeling. Scale bar: left: 50μm; right: 10μm. (**e-h**) Summary data showing GluA1CTP and GluA2CTP decreased both PSD95 (e) and gephyrin (f) puncta density in excitatory dendrites (number of dendritic segments: Control: n=250; GluA1CTP: n=170; GluA2CTP: n=210). In inhibitory dendrites, GluA2CTP, but not GluA1CTP, significantly decreased PSD95 (**g**) and gephyrin (**h**) puncta density (number of dendritic segments: Control: n=134; GluA1CTP: n=54; GluA2CTP: n=49). \**P* < 0.05; \*\**P* < 0.01; Kruskal-Wallis test with posthoc Mann-Whitney U test.

We examined the effects of GluACTP expression on the density of PSD95 and gephyrin puncta in dendrites of sparsely transfected excitatory or inhibitory tectal neurons. Excitatory and inhibitory tectal neurons receive excitatory and inhibitory synaptic inputs (Fig. 2c). Both PSD95 and gephyrin immunolabeling are highly punctate, with higher puncta density in the neuropil than the somatic region (Fig. 2d). GFP+ dendritic segments were identified as inhibitory or excitatory by GABA immunolabeling. In both excitatory and inhibitory neurons, GluA2CTP expression reduced the density of both PSD95 and gephyrin puncta (Fig. 2e-h), indicating a decrease in the number of both excitatory and inhibitory synapses onto transfected neurons. GluA1CTP expression decreased both PSD95 and gephyrin puncta in excitatory neurons but not in inhibitory neurons. Given that over 70% of tectal neurons are excitatory^19^, these results are consistent with decreased mEPSC and mIPSC frequency seen in electrophysiological recordings from randomly recorded neurons, and demonstrate cell-autonomous loss of inhibitory synaptic inputs induced by decreased excitatory input in both excitatory and inhibitory neurons.

### GluACTP expression differentially affects dendritic arbor growth and experience-dependent structural plasticity in excitatory and inhibitory neurons

To assess effects of disrupted excitatory synaptic transmission on dendritic arbor development and experience-dependent structural plasticity in excitatory and inhibitory neurons, we performed in vivo time-lapse imaging of GluACTP- and GFP-co-expressing individual neurons. Excitatory and inhibitory neurons were identified by post-hoc GABA immunolabeling (Fig. 3a-b). Total dendritic branch length (TDBL) and total branch tip number (TBTN) from 3-D reconstructions of the imaged neurons indicated that GluA1CTP and GluA2CTP expression in excitatory neurons significantly decreased TBTN but not TDBL, resulting in decreased branch density, without affecting dendritic arbor branching pattern (Fig. 3c, e). In inhibitory neurons, GluA2CTP expression significantly increased TDBL, without affecting TBTN or branch density (Fig. 3d). Interestingly, GluA2CTP expression also changed the branching pattern of inhibitory neurons, causing neurons to branch significantly farther from the soma (Fig. 3f), possibly reflecting a compensatory response to decreased excitatory inputs, consistent with observations of activity-dependent redistribution of synapse in the absence of normal activity^21^.

**Figure 3.**
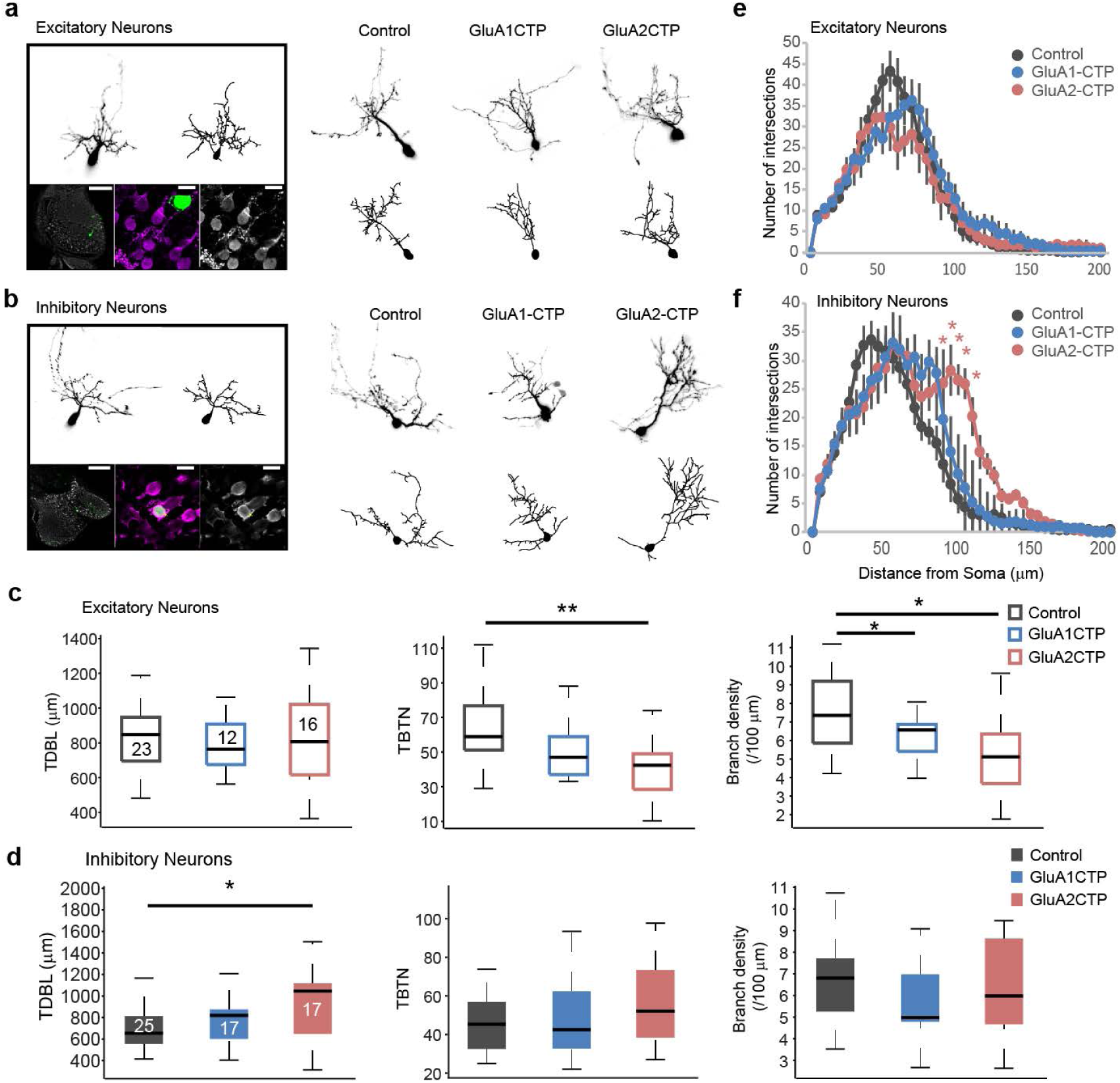
GluACTPs differentially affect dendritic arbor growth in excitatory and inhibitory neurons. (**a-b**) Representative images of excitatory (**a**) and inhibitory (**b**) neurons. Left panel: Example of the live image and reconstructed full dendritic arbor of an individual neuron (top). Post-hoc GABA immunolabeling (bottom) of the same neuron. Scale bar: left 100μm; middle and right 10μm. Right panel: Representative live images (top) of inhibitory neurons expressing GFP only (control), GluA1CTP, or GluA2CTP, and reconstructed complete dendritic arbors (bottom). (**c-d**) Summary data of total dendritic branch length (TDBL), total branch tip number (TBTN), and branch density control, GluA2CTP- and GluA1CTP-expressing excitatory (**c**) and inhibitory (**d**) neurons. *: p<0.05, Kruskal-Wallis test with posthoc Mann-Whitney U test. Number of neurons in each group is marked on the corresponding histogram bar. (**e-f**) Sholl analysis: neither GluA1CTP nor GluA2CTP affected dendritic arbor branching pattern in excitatory neurons (**e**). GluA2CTP but not GluA1CTP significantly increased branch density distal to the soma in inhibitory neurons (**f**) compared to controls. \**P* < 0.05, \*\**P* < 0.01, Kruskal-Wallis test with posthoc Mann-Whitney U test.

Previous studies showed that 4hr of short-term enhanced visual experience (STVE) increased dendritic arbor growth rate in tectal neurons compared to 4hr in dark, and that GluACTP expression blocks this visual experience-dependent dendritic arbor structural plasticity^13^. These studies imaged randomly sampled tectal neurons, therefore the results likely reflect plasticity in excitatory neurons, the majority of tectal neurons. Inhibitory tectal neurons, on the other hand, demonstrate a bimodal experience-dependent plasticity, with an inverse correlation between the valence of plasticity in response to dark and STVE in individual neurons. Furthermore, inhibitory neurons cluster into two functional groups, one group increases dendritic arbor growth rate in response to STVE and decreases arbor growth rate in the dark, while the other decreases arbor growth rate in STVE and increases it in dark.

To test if the bimodal experience-dependent structural plasticity in inhibitory neurons is affected by disrupting excitatory synaptic inputs, we collected time-lapse images of individual tectal neurons co-expressing GFP and GluA1CTP or GluA2CTP in animals exposed to 4hr of dark followed by 4hr of STVE (Fig. 4a). In excitatory neurons, GluACTP blocked the STVE-induced dendritic arbor plasticity (Control: dTDBL, p<0.01, dTBTN, p<0.05. GluA1CTP: dTDBL, p=0.52, dTBTN, p=0.83. GluA2CTP: dTDBL, p=0.94, dTBTN, p=0.82. Wilcoxon test). Comparing dendritic arbor growth rates over 4h in STVE and 4h in dark for individual neurons demonstrated that GluACTP selectively blocked the structural plasticity of excitatory neurons in response to STVE but not in dark (Fig. 4b-c). The pooled population of inhibitory neurons showed no difference in dendritic arbor plasticity between dark and STVE in controls (dTDBL, p=0.38, dTBTN, p=0.59) or GluACTP-expressing neurons (GluA1CTP: dTDBL, p=0.06, dTBTN, p=0.48. GluA2CTP: dTDBL, p=0.96, dTBTN, p=0.92, Wilcoxon test). The magnitude of structural responses to either dark or STVE was not different between GluACTP-expressing and control neurons (Fig. 4d). By contrast, plotting dendritic arbor growth rates over 4h in STVE versus dark for individual neurons demonstrates that GluACTP expression disrupted the inverse correlation between the valence of structural plasticity in dark and STVE (Fig. 4e).

**Figure 4.**
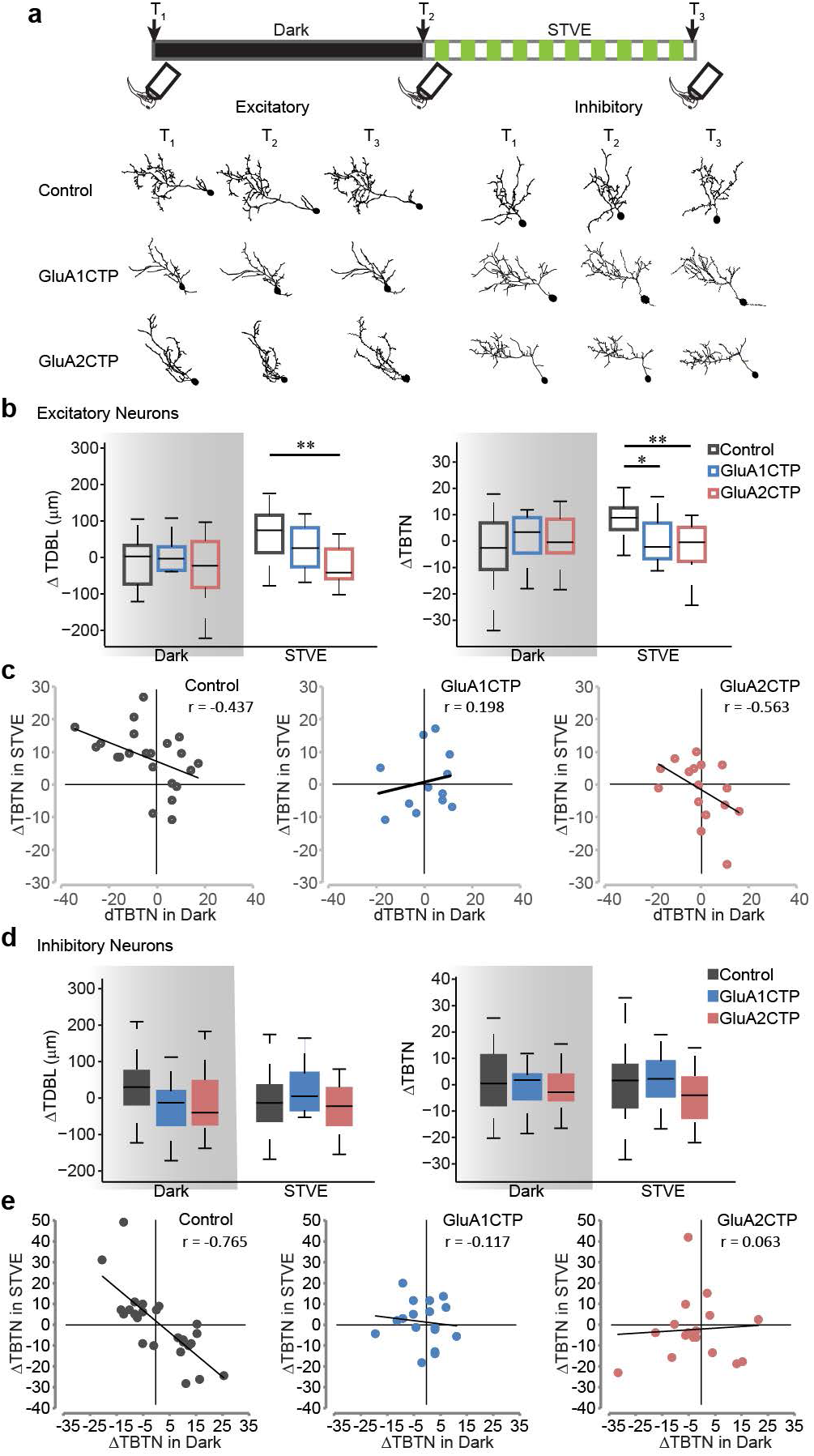
GluACTP expression disrupts experience-dependent structural plasticity in excitatory and inhibitory neurons. (**a**) Representative images of complete dendritic arbor reconstructions from time-lapse images taken before (T_1_) and after (T_2_) 4 hr of dark, and after 4 hr of STVE (T_3_) of individual excitatory and inhibitory neurons in each group. Top: schematic shows the experimental protocol and imaging time course. (**b**) Summary data of changes in TDBL and TBTN during the dark and STVE periods in excitatory neurons. GluA1CTP and GluA2CTP significantly decrease STVE-induced dendritic arbor growth. * *P* < 0.05, ** *P* < 0.01, Kruskal-Wallis test with posthoc Mann-Whitney U test. (**c**) Scatter plots of ATBTN in response to STVE versus dark in individual excitatory neurons. Rho value of Pearson correlation is shown on each plot. (**d**) Summary data. Changes in TDBL and TBTN during dark and STVE periods in inhibitory neurons. (**e**) Scatter plots of ATBTN in response to STVE vs. dark in individual inhibitory neurons.

Application of unsupervised cluster analysis based on ΔTBTN in response to STVE versus dark clustered control inhibitory neurons into two evenly-sized subpopulations, called Group I and Group II neurons (Fig. 5a, Group I: n=14; Group II: n=16). Group I and Group II neurons displayed experience-dependent structural plasticity with opposite valence, accounting for the lack of experience-dependent plasticity in the pooled population (Fig 4c). The plasticity profile of Group I inhibitory neurons was similar to excitatory neurons, retracting dendrites in the dark and extending them in STVE. The plasticity profile of Group II inhibitory neurons was the opposite, extending dendrites in the dark and retracting them in STVE (Fig. 5aiii,). We applied the same cluster analysis to GluACTP-expressing inhibitory neurons and assigned the clustered subgroups by the difference between average ΔTBTN in dark and STVE as in control neurons (Group I: ΔTBTN in STVE > ΔTBTN in dark; Group II: ΔTBTN in STVE < ΔTBTN in dark). GluA1CTP-expressing inhibitory neurons clustered into two groups with similar structural plasticity profiles as control neurons: Group I neurons retracted dendrites in dark and extended them in STVE. Group II neurons had opposite plasticity profiles. Both groups showed significant differences between plasticity in dark and STVE (Fig. 5b, Group I: n = 11; Group II: n = 6). By contrast, GluA2CTP-expressing inhibitory neurons clustered into groups that displayed distinct experience-dependent structural plasticity profiles from control neurons: Group I neurons extended dendrites and Group II neurons retracted them in both dark and STVE. Growth rates were not significantly different between dark and STVE in either subgroup (Fig. 5c, Group I: n =5, p = 0.44; Group II: n = 12, p = 0.41, Wilcoxon sign rank test), and the experience-dependent bi-modal structural plasticity in dark and STVE seen in control inhibitory neurons was abolished. In addition, comparison of the structural plasticity in response to dark or STVE between GluACTP-expressing and control inhibitory neurons showed that within both Group I and Group II neurons, GluA2CTP expression significantly altered the plasticity in response to dark, not STVE (Fig. 5d-g). By contrast, the plasticity in response to either dark or STVE in GluA1CTP-expressing inhibitory neurons was not different from controls in either group. These results indicate that GluA2 is of particular importance for the bi-modal plasticity in inhibitory neurons. Considering that only GluA2CTP-expression significantly decreased PSD95 and gephyrin puncta in inhibitory neurons (Fig. 2g-h), these results again suggest that, unlike excitatory neurons, inhibitory tectal neurons are more sensitive to the disruption of GluA2-mediated AMPAR trafficking. The loss of the bi-modal plasticity response indicates that disrupting excitatory synaptic inputs to inhibitory neurons changed their circuit connectivity^19^.

**Figure 5.**
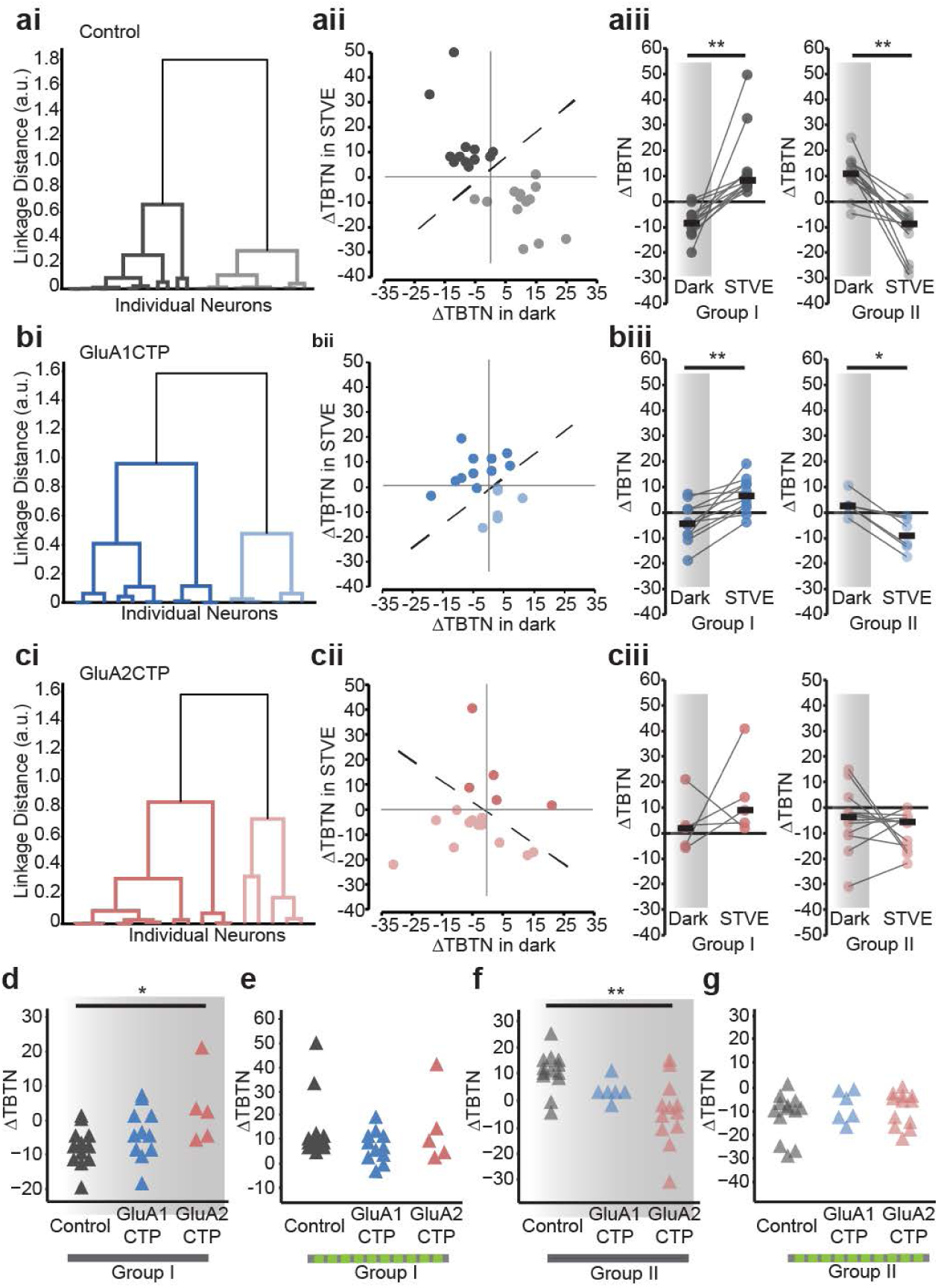
GluA2CTP expression disrupted the bimodal experience-dependent structural plasticity of inhibitory neurons. (**ai**) Dendrogram of unsupervised hierarchical cluster analysis of control inhibitory neurons based on changes in TBTN (ATBTN) in STVE versus dark. (**aii**) Scatter plots of ATBTN in response to STVE versus dark in individual neurons for Group I and II inhibitory neurons. (**aiii**) Summary of ATBTN in dark and STVE for control Group I and II neurons. (**bi-biii**) Cluster analysis of GluA1CTP-expressing inhibitory neurons. (**ci-ciii**) Cluster analysis of GluA2CTP-expressing inhibitory neurons. * *P* <0.05, ** *P* <0.01, Wilcoxon sign rank test. (**d-g**) Summary data of ΔTBTN in response to dark (**d**, **f**) and STVE (**e**, **g**) in Group I (**d**, **e**) and Group II (**f**, **g**) inhibitory neurons. * *P* <0.05, Kruskal-Wallis test with posthoc Mann-Whitney U test.

Using unsupervised cluster analysis, control excitatory neurons cluster into two groups distinguished by their plasticity in the dark (Fig. 6a, Group I: n=13; Group II: n=10): Group I excitatory neuron dendrites retract in dark and extend in STVE (n=13, p<0.001, Wilcoxon sign rank test). Group II excitatory neuron dendrites typically grow in both dark and STVE and grow significantly more in dark than in STVE (n=10, p<0.05). GluA1CTP expression significantly changed the plasticity profile in excitatory neurons: Group I neurons retract dendrites in both dark and STVE and Group II neurons extend dendrites in both dark and STVE. Responses to dark and STVE were not different within each group (Fig 6b. Group I: n=4, p=0.75. Group II: n=8, p=0.46, Wilcoxon sign rank test). Interestingly, GluA2CTP expression changed the experience-dependent structural plasticity profile of excitatory neurons to a bi-modal pattern resembling that of control inhibitory neurons: Half retract dendrites in dark and extend them in STVE, and half have the opposite plasticity profile (Fig 6c, Group I: n=8. Group II: n=8.). Comparison of the structural plasticity between GluACTP-expressing and control excitatory neurons within each group showed that GluA1CTP and GluA2-CTP significantly affected the plasticity in response to STVE but not dark (Fig. 6d-g). The observation that decreased excitatory inputs significantly affected the STVE response in excitatory neurons and the dark response in inhibitory neurons provides further evidence that the plasticity of inhibitory tectal neurons is actively regulated in dark^19,22^.

**Figure 6.**
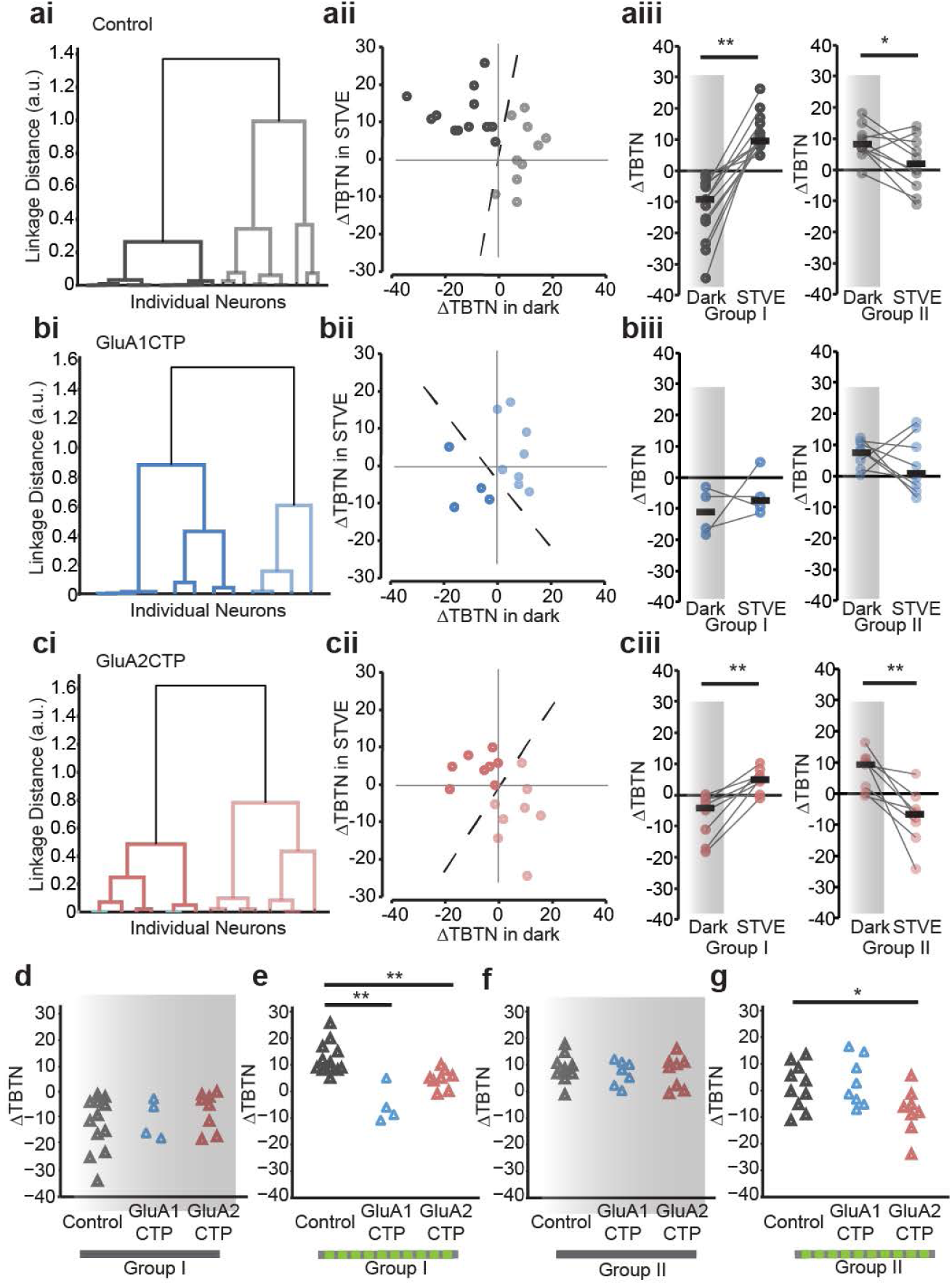
Cluster analysis of experience-dependent TBTN changes of individual excitatory neurons. (**ai**) Dendrogram of unsupervised hierarchical cluster analysis of control excitatory neurons based on dTBTN in STVE versus dark. (**aii**) Scatter plots of ATBTN in response to STVE vs dark in individual neurons for Group I and II excitatory neurons. (**aiii**) Summary of ATBTN in dark and STVE for Group I and II control neurons. (**bi-biii**) Cluster analysis of GluA1CTP-expressing excitatory neurons. (**ci-ciii**) Cluster analysis of GluA2CTP-expressing excitatory neurons. * *P* <0.05, ** *P* <0.01, Wilcoxon sign rank test. (**d-g**) Summary data of ATBTN in response to dark (**d, f**) and STVE (**e, g**) in group I (**d**, **e**) and group II (**f, g**) excitatory neurons. \**P* <0.05, Kruskal-Wallis test with posthoc Mann-Whitney U test.

### Decreased excitatory and inhibitory inputs disrupt receptive field properties

Interaction of excitatory and inhibitory inputs is thought to be essential for the developmental refinement of visual receptive field (RF) properties^23–25^. Here, we showed that interfering with glutamatergic inputs compromises both excitatory and inhibitory inputs yet E/I remains constant. To test whether RF properties are affected by GluACTP expression, we measured the spatial and temporal RF properties in transfected neurons. We recorded spatial receptive fields in both cell-attached mode, to measure the spiking receptive fields (sRF), and whole-cell mode, to measure excitatory receptive fields (eRF) and inhibitory receptive fields (iRF) respectively in response to light off visual stimuli (Fig. 7a). sRFs, eRFs and iRFs were all significantly smaller in GluA1CTP- and GluA2CTP-expressing neurons than controls (Fig. 7b-d). GluACTP expression disrupted the convergence of eRF and iRF as shown by significantly greater distance between the eRFs and iRFs centers (Fig. 7e). We generated temporal receptive field (tRF) maps by binning the number of spikes in 100 ms intervals over the 700 ms recording period following the OFF stimulus (Fig. 7f). The tRFs in control GFP+ neurons was confined to 200 ms following the stimulus (Fig. 7g). By contrast, the tRF in GluA1CTP- and GluA2CTP-expressing neurons extended up to 600 ms after the stimulus. Consequently, the average spike latency and the full width at the half maximal response (FWHM) of the tRFs in GluA1CTP- and GluA2CTP-expressing neurons were significantly greater than controls (Fig. 7h-i). The decreased convergence of eRF and iRF and the increased temporal span of the visually evoked spikes are consistent with a decreased inhibition^15,18,24,25^. Together, these data indicate that decreasing excitatory synaptic inputs and the subsequent cell autonomous decrease in inhibitory synaptic inputs in tectal neurons disrupted the tectal circuits underlying visual information processing and impaired both spatial and temporal RF properties.

**Figure 7.**
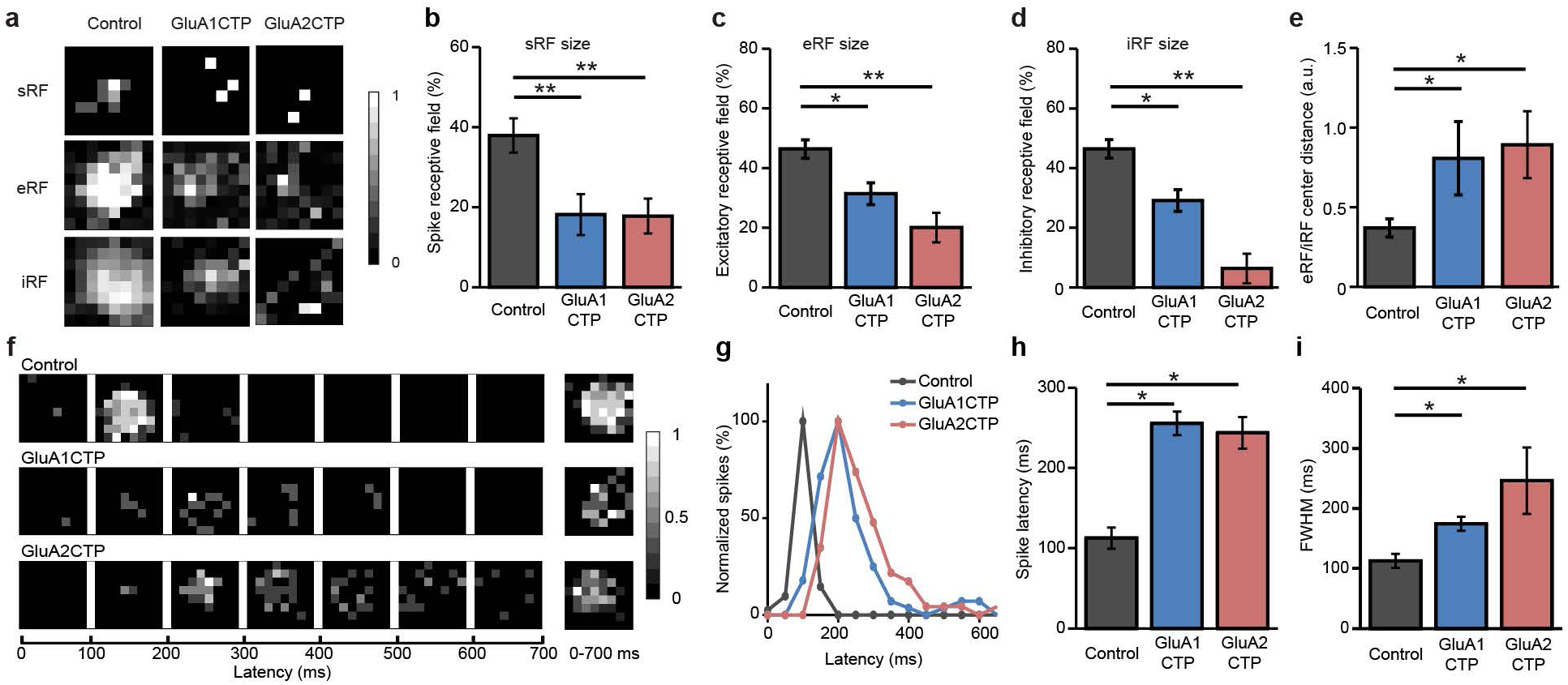
GluACTP expression disrupts spatial and temporal receptive visual field properties. (**a**) Representative maps of spiking RFs (sRFs), excitatory RFs (eRFs), and inhibitory RFs (iRFs) in control, GluA1CTP - and GluA2CTP-expressing neurons. (**b-d**) Both GluA1CTP and GluA2CTP expression significantly decreased the size of sRF (**b**, n=28, 14, 8), eRF (**c**, n=18, 19, 20), and iRF (**d**, n=18, 12, 13). (**e**) The distances between eRF and iRF centers were significantly larger in GluA1CTP and GluA2CTP-expressing neurons. (**f**) Representative tRF maps in control, GluA1CTP- and GluA2CTP-expressing neurons. (**g**) Plot of spike numbers over the 700 ms recording period normalized to peak spike numbers per 100 ms bin. The control tRF is limited to the first 200 ms while tRFs in GluA1CTP- and GluA2CTP-expressing neurons were significantly extended and delayed. (**h-i**) Both spike latency (**h**) and the full width at half maximum (FWHM, **i**) of the tRF spiking response increased significantly in GluA1CTP- and GluA2CTP-expressing neurons compared to control neurons. Control: n=12; GluA1CTP: n=9; GluA2CTP: n=8. * *P* <0.05, ** *P* <0.01. ANOVA with Newman-Keuls test.

### GluACTP expression impairs visual avoidance behavior and learning-induced behavioral plasticity

We next addressed whether GluACTP expression affects visual avoidance behavior and behavioral plasticity. The results described above predict two outcomes of GluACTP expression: On one hand, if constant E/I is sufficient for circuit function underlying behavior, then the co-regulation of inhibitory inputs in response to GluACTP expression and the resultant constant E/I, predicts that visual avoidance behavior would be intact. By contrast, GluACTP expression impaired visual information processing and RF properties, predicting that circuit function and plasticity would be impaired. We bulk electroporated tectal neurons with GluA1CTP, GluA2CTP, or GFP expression constructs and evaluated visual avoidance behavior, and avoidance behavior plasticity. The visual avoidance behavior is a tectal-mediated visually-guided behavior in which an animal changes swim trajectory in response to an approaching visual stimulus^15^. Animals improve their avoidance behavior following a visual-conditioning training protocol^26^. Control tadpoles avoided dots with diameters ranging from 0.2 to 0.6 cm, with the peak avoidance response to dots of 0.4 cm diameters (Fig. 8a-b). GluA1CTP- and GluA2CTP-expressing tadpoles had significantly lower avoidance indices to stimuli of 0.4 cm and 0.2 cm, and GluA2CTP-expressing animals also showed a significantly decreased response to 0.6 cm dots (Fig. 8b), indicating impaired visual behavior in these animals despite of balanced E/I. GluA1CTP and GluA2CTP expression also blocked visual-conditioning mediated plasticity of the behavior (Fig. 8c). This learning deficit is consistent with the compromised experience-dependent structural plasticity and visual information processing observed in individual neurons expressing GluACTPs.

**Figure 8.**
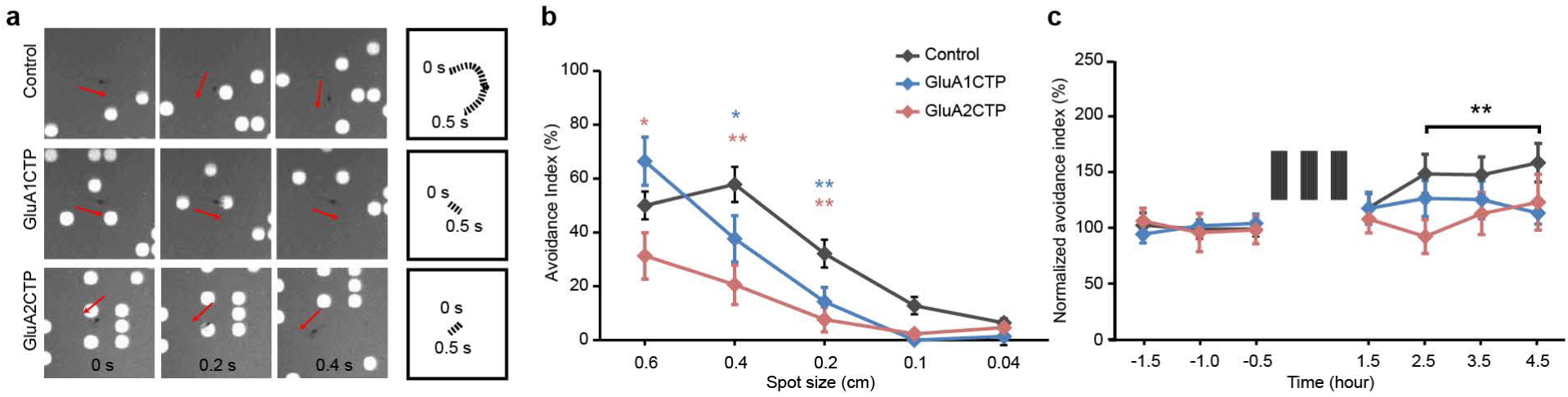
GluACTP expression impairs visual avoidance behavior and behavioral plasticity. (**a**) Representative snapshots of tadpole behavior in response to upward moving spot stimuli (diameter 0.4 cm) in animals expressing GFP, GluA1CTP or GluA2CTP in tectal neurons. Top panel: Control tadpoles turned to avoid an approaching stimulus. The swim trajectory over 500ms is shown on the right. Tadpoles in which the optic tectum was electroporated with GluA1CTP (middle panel) or GluA2CTP (bottom panel) did not change swim trajectories in response to a moving stimulus. (**b**) Summary data: Avoidance index in response to stimuli of diameters 0.04–0.6 cm for animals expressing GFP (control, n=15) or GFP with GluA1CTP (n=21) or GluA2CTP (n=18) in the optic tectum. (**c**) GluA1CTP (n=20) or GluA2CTP (n=16) expression blocked visual experience-induced enhancement of visual avoidance behavior observed control tadpoles (n=18). * *P* <0.05, ** *P* <0.01. ANOVA with Newman-Keuls test.

## Discussion

Genetic variants of proteins associated with glutamatergic synaptic function, such as CTNAP2 and SHANK3, have been implicated in the etiology of neuropsychiatric disorders, placing excitatory synapse dysfunction in the spotlight as a candidate mechanism underlying pathogenesis of these disorders^27,28^. Animal models with these genetic deficits also show reduced inhibitory tone, recapitulating human patient studies^29^. Some of these neurological disorders are thought to have neurodevelopmental origins, such as ASD and schizophrenia, raising the intriguing questions: Is there a causal link between defective excitatory synaptic function during development and reduced inhibition? And how do deficits in excitatory synaptic functions lead to deficits in inhibitory function?

Here we examined the role of glutamatergic excitatory synaptic transmission in the coordinated development of inhibition and excitation at synaptic, neuronal, circuit and behavioral levels, focusing on inhibitory synapses and neurons. We report a coordinated cell-autonomous reduction in synaptic inhibition in response to decreasing glutamatergic transmission by GluACTP expression in individual neurons. The decreased inhibitory input was not only observed in spontaneous activity but also in visually evoked inhibitory synaptic responses, which drastically altered the spatial and temporal visual receptive field properties in transfected tectal neurons. Disrupting excitatory transmission also blocked experience-dependent structural plasticity in both inhibitory and excitatory neurons. Interestingly, the structural plasticity deficit in inhibitory neurons occurred in the response to dark, whereas in excitatory neurons the deficit occurred in response to visual stimulation. These synaptic and cellular defects translated into behavioral deficits when GluACTPs were expressed more extensively in tectal neurons.

The C-terminals of AMPARs include core regulatory sites for AMPAR trafficking. CTPs compete with endogenous AMPARs for binding partners, thereby interfering with AMPAR trafficking into synapses and decreasing excitatory synaptic transmission. Different mechanisms may underlie the disruption of excitatory synaptic inputs by expression of GluA1CTP and GluA2CTP due to different synaptic delivery mechanisms of GluA1- and GluA2-containing AMPARs^11,12^. GluA1CTP does not affect basal AMPAR-mediated currents but abolishes activity-dependent synaptic potentiation. GluA2CTP significantly decreases basal AMPAR-mediated synaptic transmission, which produces greater synaptic potentiation in response to LTP-inducing protocols, but due to impaired synaptic delivery of GluA2-containing receptors, this increased synaptic strength is not maintained^9^. AMPAR trafficking also affects homeostatic plasticity where GluACTP blocks inactivation-induced synaptic scaling^30^. Nevertheless, expression of either GluA1CTP or GluA2CTP compromises excitatory synaptic transmission, due to impaired basal transmission and synaptic plasticity.

Although many studies have examined circuit-wide regulation of E/I balance^2,4,6,31^, little is known about whether cell autonomous mechanisms play a role in this critical aspect of neuronal and circuit function. Knocking down glutamatergic transmission in a small subset of neurons reduced both mIPSC and evoked IPSCs in hippocampal slice culture, suggesting cell-autonomous regulation of E/I^32^. Our results showing the proportional decrease in excitatory and inhibitory inputs in sparsely-transfected GluACTP-expressing neurons provides in vivo evidence that cell-autonomous mechanisms maintain E/I. By contrast, we previously reported that decreasing inhibition by expressing a peptide interfering with GABAA receptor trafficking does not affect glutamatergic synaptic inputs onto the same neuron, thereby disrupting E/I and causing dysfunction of the tectal circuit^15^. These results suggest that cell-autonomous modification of E/I is triggered by a direct change in excitatory synaptic input and not by the net change of excitatory inputs relative to inhibitory inputs.

The cell-autonomous downregulation of inhibition following disruption of excitatory inputs may have important implications for the etiology of some neurological disorders. Loss of function of the autism-related cell adhesion molecule CNTNAP2 in cultured cortical neurons caused a cell-autonomous decrease in both excitatory and inhibitory synaptic input^33^. CNTNAP2 preferentially co-localizes with GluA1 and knocking it down in cultured neurons led to abnormal cytoplasmic aggregation of GluA1, suggesting a role in AMPAR trafficking^34^. The dysfunction of inhibitory synaptic transmission observed with CNTNAP2 knockdown may be a secondary consequence of disrupted excitatory transmission due to defective AMPAR trafficking, as we observed here. Another consequence of the coordinated decrease in inhibition induced by decreased excitation is that E/I remained relatively stable. E/I is thought to be critical for neural circuit stability and normal brain function^2^. Disrupted E/I is associated with several neurological diseases, including epilepsy, schizophrenia and autism spectrum disorders^35–37^. Here we showed that even though E/I was resilient to disruption of excitatory inputs, nervous system function was still significantly compromised at both neuronal and circuit levels, shown by abnormal dendritic morphology, altered experience-dependent plasticity and defective receptive field properties, suggesting that balanced E/I is not sufficient to maintain normal brain function. Mice with MECP2 knockout in forebrain excitatory neurons provide another example of simultaneous reduction in excitation and inhibition resulting in functional deficits despite relatively balanced E/I^38^.

Classical studies have shown that excitatory sensory inputs modify inhibition, however studies specifically investigating the direct cell autonomous involvement of AMPAR-mediated excitatory inputs in the development of inhibitory neurons have been sparse^39^. Visual deprivation in both young and adult rodent decreases inhibition in primary visual cortex^40,41^. Monocular deprivation decreases inhibition by increasing turnover and net loss of inhibitory synapses in adult visual cortex^42,43^. Interfering with AMPAR-mediated inputs specifically in parvalbumin neurons, either with GluA1 or GluA4 knockout, or by manipulating neuronal pentraxins, delayed inhibitory circuit maturation, blocked cortical plasticity and resulted in behavioral deficits^44–46^. Visually-induced potentiation and depression of GABAergic synapses have been shown to be correlated with the E/I of postsynaptic cells in the optic tectum^47^. These prior studies did not distinguish between circuit and cell-autonomous control of inhibition. Here, with in vivo manipulation of AMPAR-mediated synaptic inputs in single neurons, we showed that disrupting excitatory inputs onto developing neurons resulted in coordinated decrease in both excitatory and inhibitory synaptic inputs, indicating that both excitatory and inhibitory neurons are subject to cell-autonomous regulation of inhibitory inputs by excitatory inputs.

Specific cellular mechanisms underlying the cell-autonomous down regulation of inhibition by decreased excitatory inputs are still unclear. One potential mechanism is retrograde signaling through BDNF, which has been shown to regulate formation and maintenance of inhibitory inputs in response to excitatory inputs^48^. NO has also been proposed as a retrograde messenger that mediates heterosynaptic potentiation of GABAergic synapses^49^. Previous studies indicate that cell-autonomous regulation of inhibitory synaptic inputs is independent of postsynaptic spiking^4^, suggesting that excitatory postsynaptic activity alone is sufficient to control the formation and maintenance of inhibitory synapses. When GABAergic currents are hyperpolarizing and AMPARs are major source of synaptic depolarization, as is the case in our experiments, disrupting AMPAR-mediated excitatory synaptic inputs reduces the depolarization that is crucial for the activation of NMDARs, which has been shown to be obligatory for the development of inhibitory synaptic transmission^49,50^. Our data provide direct evidence for an essential role for glutamatergic excitatory transmission in the cell autonomous development of GABAergic inhibition and distinguished effects of excitatory inputs on excitatory and inhibitory neurons.

## Materials and methods

### Animals and Transfection

Albino Xenopus laevis were reared as previously described^19^. All animal protocols were approved by the Institutional Animal Care and Use Committee of the Scripps Research Institute and the local ethics committee of the Hangzhou Normal University. Stage 46–48 animals were anesthetized in 0.02% MS-222 (Tricane methanesulfonate, Sigma, St. Louis, MO) and the tectum was co-electroporated with pGal4 and either UAS::GluA1CTP-T2A-GFP or UAS::GluA2CTP - T2A- GFP.

### Electrophysiology

All recordings were performed at room temperature (20 - 22°C). During the recordings, brains were perfused with extracellular saline containing (in mM: 115 NaCl, 2 KCl, 3 CaCl2, 1.5 MgCl2, 5 HEPES, 10 glucose, 0.01 glycine and 0.05 Tubocurarine, pH 7.2, osmolality 255 mOsm). Visually evoked synaptic currents were recorded from tectal neurons in the middle of the tectum in whole cell mode using a K+-based pipette solution (in mM: 110 K-gluconate, 8 KCl, 5 NaCl, 1. 5 MgCl_2_, 20 HEPES, 0.5 EGTA, 2 ATP, and 0.3 GTP). Action potentials were recorded in cell attached mode. Recording micropipettes were pulled from borosilicate glass capillaries and had resistances in the range of 7 - 9 MΩ. Liquid junction potential was adjusted during recording. Whole cell recordings were accepted for analysis from cells in which the series resistance did not change over 10% and input resistance (0.7 - 2 GΩ) remained relatively constant. Signals were filtered at 2 kHz with a Multiclamp 700A amplifier (Molecular Devices, Palo Alto, CA). Data were sampled at 10 kHz using ClampFit 10 (Molecular Devices). Responses, including spikes, excitatory compound synaptic currents (eCSCs) and inhibitory compound synaptic currents (iCSCs) to light off stimuli were analyzed by Matlab (The MathWorks, Psychophysics Toolbox extensions).

For whole-cell recordings, tadpoles were anesthetized and tectal lobes were cut along the dorsal midline with a sharp needle. Live tadpoles were immobilized on a sylgard cushion in the recording chamber with one eye facing the center of a back-projection screen. Full field visual stimuli were generated in Matlab with Physics Toolbox and presented from lowest to highest luminance (10, 20 and 250 cd/m^2^) from a projector (Samsung, sp-p310ME LED projector) to the back-projection screen. Each stimulus was presented 10 times (Frequency: 0.1 Hz, interval: 0.05 Hz).

For receptive field mapping, white squares on a dark background were presented in an 8×8 grid of 0.5×0.5 cm^2^ non-overlapping squares covering a 4×4 cm^2^ area in the visual field. The entire visual field was mapped by randomly presenting white squares for 1.5 s with 5 s intervals. For spike receptive field mapping, using cell attached recordings, the response within 700 ms after the onset of the off stimulus at each grid position was determined as the average number of total spikes per stimulus from two or three repeats. After cell-attached recording, whole cell voltage clamp recording was accomplished by applying negative pressure. The same visual stimuli were used to measure inhibitory spatial receptive fields and excitatory spatial receptive fields, by holding membrane potential at 0 mV and −60 mV respectively. Total synaptic charge transfer over 700 ms from the onset of stimulus was normalized to the peak response and computed by Matlab to show spatial receptive field size. All values larger than 3 times the standard deviation of spontaneous activity were included in the measurements of spatial receptive fields. The center of the spatial receptive field was defined as the center of the square that elicited the highest responses (maximum number of spikes).

### In Vivo Time Lapse Imaging of Dendritic Arbor Structure and Data Analysis

Animals were electroporated with DNA constructs at stage of 45–46 and screened for those with sparsely transfected and well-isolated cells. For imaging, animals were anesthetized with 0.01% MS-222 (Sigma) and were placed in a Sylgard chamber covered by a glass coverslip. Images were collected every 4hr before and after each visual experience session (dark or STVE). Two-photon z-series were collected at 1 μm steps with a 20x water immersion objective (Olympus XLUMPlanFL 0.95NA) at 3–4x scan zoom using a custom-built microscope modified from an Olympus FV300 system^19^.

Complete dendritic arbors of each neuron were reconstructed using a semi-manual function in the Filament module of Imaris (Bitplane, US). Total dendritic length and branch tip number were automatically calculated by the software. 3D Sholl analysis calculated the number of branches that intersect concentric circles at increasing distances from the cell soma, using a customized Matlab program with reconstructed filament data exported from Imaris.

### Immunohistochemistry and image data analysis

For posthoc analysis of neurons imaged for in vivo time-lapse experiments, animals were fixed at the end of the imaging experiment with freshly made 4% paraformaldehyde and 2% Glutaraldehyde (Electron Microscopy Sciences, Fort Washington, PA) in 1xPBS (pH 7.4) using a Pelco BioWave Pro microwave (Model 36500, Ted Pella, Redding, CA. 350mV on 20 sec, off 20 sec, on 20 sec, followed by 150mV on 1 min, off 1 min, on 1 min). The animals were then post-fixed at 4°C overnight and washed in 1xPBS using the microwave (150mV on-off-on, 1min each). 30μm vibratome sections of the dissected brains were cut for free floating immunofluorescence labeling. Sections were incubated in 1% Sodium Borohydride (Sigma) in 1xPBS for 15 min to quench auto fluorescence, blocked in 10% normal goat serum (Jackson Lab, ME) in PBS with 2% Triton X-100 (PBST) for one hour in room temperature, followed by incubation in rabbit anti GABA polyclonal antibody (Sigma A2052, RRID: AB_477652, 1:2000 in PBST with 1% normal goat serum) for 48 hours at 4°C. Secondary antibody (goat anti rabbit Alexa Fluor 633, Thermo Fisher Scientific Cat# A-21070, RRID: AB_2535731) was diluted 1:500 in PBST and incubated for an hour at room temperature. After 3x 15min rinses with PBS, sections were mounted on slides in Vectashield Mounting Medium with DAPI (Vector Laboratories, Burlingame, CA). For immunolabeling with GluA antibodies, stage 47 animals were fixed in 4% PFA and 0.5% glutaraldehyde. Primary antibodies used include mouse anti GluA2 (N) antibody (Millipore Cat# MAB397, RRID: AB_2113875) and mouse anti GluA1-N antibody (generous gifts from Dr. R. Huganir, Johns Hopkins University Medical School). For PSD95 and gephyrin puncta analysis, 7 days after electroporation of the corresponding DNA constructs, animals were fixed with 4% PFA with 1 hour post-fix at room temperature. Free floating vibratome sections were quenched and blocked as mentioned above, followed by incubation in the primary antibodies including rabbit-anti-PSD95 polyclonal antibody (1:50, Thermo Fisher Scientific Cat#51–6900, RRID: AB_2533914), mouse-anti-gephyrin monoclonal antibody (1:50, Synaptic Systems Cat#147021, RRID: AB_2232546), guinea pig-anti-GABA polyclonal antibody (1:1500, Millipore Cat# AB175, RRID: AB_91011). Secondary antibodies used include goat-anti-guinea pig Alexa Fluor 568 (Thermo Fisher Scientific Cat# A-11075, RRID: AB_141954), goat-anti-rabbit BV421 (BD Biosciences Pharmingen, Cat#565014, RRID: AB_2716308), donkey-anti-mouse Alexa Fluor 647 (Thermo Fisher Scientific Cat# A-31571, RRID: AB_162542). Brains from control and experimental groups were embedded in the same blocks and processed under exactly same conditions throughout the experiments.

Images of immunolabeled sections for posthoc identification of GABAergic neurons for the in vivo time-lapse imaging experiments were collected on an Olympus Fluoview 500 confocal microscope or a Nikon C2 confocal microscope. GFP fluorescence signal was well preserved in the fixed tissue, thus no antibody was needed for visualization. The GFP+ imaged neurons were located using a 20x air objective and confirmed by location and dendritic arbor branching patterns. Higher magnification images were then taken at > 3 different z-depth through the soma to examine GABA immunoreactivity. Samples with poor GABA immunostaining were not included in the analysis.

Images of PSD95 and Gephyrin immunolabeling were acquired on a Nikon C2 confocal microscope with a 40x PlanFluor Oil objective (N.A. 1.3) at 2048×2048 to achieve a final resolution of 0.15 μm/pixel. Analysis of dendritic puncta density was performed using SynPAnal program^31^. Transfected dendritic segments of control or experimental groups were defined solely based on the GFP signal. The dendritic segment was deemed inhibitory if the average GABA immunolabeling intensity within the segment was above a threshold set by mean+2SD of GABA immunolabeling intensity of 3 randomly drawn GFP-segments of similar length and width in the neuropil region within the same section adjacent to the transfected dendritic segment. In a subset of dendritic segments (n=496), for which the soma could be identified, 82.5% showed GABA immunoreactivity consistent with their soma, confirming the reliability of identifying GABAergic inhibitory dendritic segments based on GABA immunoreactivity within the segments. Puncta density values (per unit length of dendrite) of PSD95 and Gephyrin puncta within the transfected dendritic segments were automatically taken by the program and were normalized to the average puncta density of the neuropil regions within the same section to control for immunostaining variability across sections. All image analysis was done blind to the treatment.

### Visual Avoidance Assay and Visual Conditioning

The visual avoidance assay was conducted as previously described^15^. Tadpoles were placed in an 8X3 cm tank filled to a depth of ~1 cm with Steinberg’s rearing solution. Visual stimuli were presented to a back-projection screen on the bottom of the chamber using a microprojector (3M, MPro110). Videos of tadpoles illuminated by IR LEDs were recorded with a Hamamatsu ORCA-ER digital camera. Visual stimuli were generated and presented by MATLAB. Randomly positioned moving spots of 0.04, 0.2, 0.4 and 0.6 cm diameter were presented in random order for 60 seconds. Visual avoidance behavior was scored as a change in the swimming trajectory in the first ten encounters between each tadpole and moving spots (the percentage of avoidance responses out of 10 encounters, plotted as avoidance index). For visual conditioning, animals were exposed to moving bars (1 cm width; 0.3 Hz; Luminance: 25 cd/m^2^) continuously for 2 or 4 hours, or for 3 repeats of 5 minutes of moving bars interleaved by 5-minute blank, for a total of 30 minutes, as described^26^.

### Cluster Analysis and Statistical Tests

Custer analysis was performed based on ΔTBTN over STVE versus darl of individual neurons using an unsupervised agglomerative hierarchical tree method in MATLAB (linkage.m) based upon their pair-wise vectorial distance in the constructed 2D space (pdist.m)^19^.

All data are presented as mean ± SEM. Data are considered significantly different when p values are less than 0.05. Where noted, either two-tailed Student’s t-test or nonparametric Wilcoxon sign rank test was performed for within-cell comparison. For comparisons of multiple groups, either ANOVA with Newman-Keuls test or Kruskal-Wallis test with posthoc Mann-Whitney U test were performed. The statistical test used for each experiment is specified in the results.

Experiments and analysis were performed blind to the experimental conditions.

## Acknowledgements

We thank Dr. Richard Huganir for sharing the GluA1 antibody and Dr. Eric Danielson for providing the SynPAnal puncta analysis software. We thank Dr. Daniel Choquet, Dr. Anton Maximov, Dr. Richard Huganir and members of the Cline lab for critical discussions. This work was supported by the NEI (EY011261 and EY 027437) and the Hahn Family Foundation to HTC, and National Nature Sciences Foundation of China (NSFC 31271176) and Zhejiang Provincial Natural Science Foundation of China (LY17C090007) to WS.

## References

1 Katz, L. C. & Shatz, C. J. Synaptic activity and the construction of cortical circuits. Science 274, 1133–1138 (1996).

2 Froemke, R. C. Plasticity of cortical excitatory-inhibitory balance. Annu Rev Neurosci 38, 195–219, doi:10.1146/annurev-neuro-071714-034002 (2015).

3 Wamsley, B. & Fishell, G. Genetic and activity-dependent mechanisms underlying interneuron diversity. Nat Rev Neurosci 18, 299–309, doi:10.1038/nrn.2017.30 (2017). 21

4 Hartman, K. N., Pal, S. K., Burrone, J. & Murthy, V. N. Activity-dependent regulation of inhibitory synaptic transmission in hippocampal neurons. Nat Neurosci 9, 642–649, doi:nn1677 [pii] 10.1038/nn1677 (2006).

5 Huang, Z. J., Di Cristo, G. & Ango, F. Development of GABA innervation in the cerebral and cerebellar cortices. Nat Rev Neurosci 8, 673–686, doi:10.1038/nrn2188 (2007).

6 Jiang, B., Huang, Z. J., Morales, B. & Kirkwood, A. Maturation of GABAergic transmission and the timing of plasticity in visual cortex. Brain Res Brain Res Rev 50, 126–133, doi:10.1016/j.brainresrev.2005.05.007 (2005).

7 Pieraut, S. et al. Experience-dependent remodeling of basket cell networks in the dentate gyrus. Neuron 84, 107–122, doi:10.1016/j.neuron.2014.09.012 (2014).

8 Bredt, D. S., & Nicoll, R. A. AMPA receptor trafficking at excitatory synapses. Neuron 40, 361–379 (2003).

9 Kessels, H. W., & Malinow, R. Synaptic AMPA receptor plasticity and behavior. Neuron 61, 340–350, doi:10.1016/j.neuron.2009.01.015 (2009).

10 Wu, G., Malinow, R. & Cline, H. T. Maturation of a central glutamatergic synapse. Science 274, 972–976 (1996).

11 Zhou, Z. et al. The C-terminal tails of endogenous GluA1 and GluA2 differentially contribute to hippocampal synaptic plasticity and learning. Nat Neurosci, doi:10.1038/s41593-017-0030-z (2017).

12 Shi, S., Hayashi, Y., Esteban, J. A. & Malinow, R. Subunit-specific rules governing AMPA receptor trafficking to synapses in hippocampal pyramidal neurons. Cell 105, 331–343 (2001).

13 Haas, K., Li, J. & Cline, H. T. AMPA receptors regulate experience-dependent dendritic arbor growth in vivo. Proc Natl Acad Sci U S A 103, 12127–12131, doi:10.1073/pnas.0602670103 (2006).

14 Yoon, B. J., Smith, G. B., Heynen, A. J., Neve, R. L. & Bear, M. F. Essential role for a long-term depression mechanism in ocular dominance plasticity. Proc Natl Acad Sci U S A 106, 9860–9865, doi:10.1073/pnas.0901305106 (2009).

15 Shen, W., McKeown, C. R., Demas, J. A. & Cline, H. T. Inhibition to excitation ratio regulates visual system responses and behavior in vivo. J Neurophysiol 106, 2285–2302, doi:10.1152/jn.00641.2011 (2011).

16 Rumpel, S., LeDoux, J., Zador, A. & Malinow, R. Postsynaptic receptor trafficking underlying a form of associative learning. Science 308, 83–88 (2005).

17 Akerman, C. J. & Cline, H. T. Refining the roles of GABAergic signaling during neural circuit formation. Trends in neurosciences 30, 382–389, doi:10.1016/j.tins.2007.06.002 (2007).

18 Khakhalin, A. S., Koren, D., Gu, J., Xu, H. & Aizenman, C. D. Excitation and inhibition in recurrent networks mediate collision avoidance in Xenopus tadpoles. Eur J Neurosci 40, 2948–2962, doi:10.1111/ejn.12664 (2014).

19 He, H. Y., Shen, W., Hiramoto, M. & Cline, H. T. Experience-Dependent Bimodal Plasticity of Inhibitory Neurons in Early Development. Neuron 90, 1203–1214, doi:10.1016/j.neuron.2016.04.044 (2016).

20 Aizenman, C. D., Munoz-Elias, G. & Cline, H. T. Visually driven modulation of glutamatergic synaptic transmission is mediated by the regulation of intracellular polyamines. Neuron 34, 623–634 (2002).

21 Chen, Y. et al. Activity-induced Nr4a1 regulates spine density and distribution pattern of excitatory synapses in pyramidal neurons. Neuron 83, 431–443, doi:10.1016/j.neuron.2014.05.027 (2014).

22 Gambrill, A. C., Faulkner, R. & Cline, H. T. Experience-dependent plasticity of excitatory and inhibitory intertectal inputs in Xenopus tadpoles. J Neurophysiol, jn 00611 02016, doi:10.1152/jn.00611.2016 (2016).

23 Carvalho, T. P. & Buonomano, D. V. Differential effects of excitatory and inhibitory plasticity on synaptically driven neuronal input-output functions. Neuron 61, 774–785, doi:10.1016/j.neuron.2009.01.013 (2009).

24 Richards, B. A., Voss, O. P. & Akerman, C. J. GABAergic circuits control stimulus-instructed receptive field development in the optic tectum. Nat Neurosci 13, 1098–1106, doi:10.1038/nn.2612 (2010).

25 Tao, H. W. & Poo, M. M. Activity-dependent matching of excitatory and inhibitory inputs during refinement of visual receptive fields. Neuron 45, 829–836, doi:10.1016/j.neuron.2005.01.046 (2005).

26 Shen, W. et al. Acute synthesis of CPEB is required for plasticity of visual avoidance behavior in Xenopus. Cell reports 6, 737–747, doi:10.1016/j.celrep.2014.01.024 (2014).

27 Monteiro, P. & Feng, G. SHANK proteins: roles at the synapse and in autism spectrum disorder. Nat Rev Neurosci 18, 147–157, doi:10.1038/nrn.2016.183 (2017).

28 Volk, L., Chiu, S. L., Sharma, K. & Huganir, R. L. Glutamate synapses in human cognitive disorders. Annu Rev Neurosci 38, 127–149, doi:10.1146/annurev-neuro-071714-033821 (2015).

29 Rapanelli, M., Frick, L. R. & Pittenger, C. The Role of Interneurons in Autism and Tourette Syndrome. Trends in neurosciences 40, 397–407, doi:10.1016/j.tins.2017.05.004 (2017).

30 Gainey, M. A., Hurvitz-Wolff, J. R., Lambo, M. E. & Turrigiano, G. G. Synaptic scaling requires the GluR2 subunit of the AMPA receptor. J Neurosci 29, 6479–6489, doi:10.1523/JNEUROSCI.3753-08.2009 (2009).

31 Danielson, E. & Lee, S. H. SynPAnal: software for rapid quantification of the density and intensity of protein puncta from fluorescence microscopy images of neurons. PloS one 9, e115298, doi:10.1371/journal.pone.0115298 (2014).

32 Lu, W., Bushong, Eric A., Shih, Tiffany P., Ellisman, Mark H. & Nicoll, Roger A. The Cell-Autonomous Role of Excitatory Synaptic Transmission in the Regulation of Neuronal Structure and Function. Neuron 78, 433–439, doi:http://arxiv.org/abs/dx.doi.org/10.1016/j.neuron.2013.02.030 (2013).

33 Anderson, G. R. et al. Candidate autism gene screen identifies critical role for cell-adhesion molecule CASPR2 in dendritic arborization and spine development. Proc Natl Acad Sci U S A 109, 18120–18125, doi:10.1073/pnas.1216398109 (2012).

34 Varea, O. et al. Synaptic abnormalities and cytoplasmic glutamate receptor aggregates in contactin associated protein-like 2/Caspr2 knockout neurons. Proc Natl Acad Sci U S A 112, 6176–6181, doi:10.1073/pnas.1423205112 (2015).

35 Fritschy, J. M. Epilepsy, E/I Balance and GABA(A) Receptor Plasticity. Front Mol Neurosci 1, 5, doi:10.3389/neuro.02.005.2008 (2008).

36 Lewis, D. A., Hashimoto, T. & Volk, D. W. Cortical inhibitory neurons and schizophrenia. Nat Rev Neurosci 6, 312–324, doi:10.1038/nrn1648 (2005).

37 Nelson, S. B., & Valakh, V. Excitatory/Inhibitory Balance and Circuit Homeostasis in Autism Spectrum Disorders. Neuron 87, 684–698, doi:10.1016/j.neuron.2015.07.033 (2015).

38 Banerjee, A. et al. Jointly reduced inhibition and excitation underlies circuit-wide changes in cortical processing in Rett syndrome. Proc Natl Acad Sci U S A 113, E7287–E7296, doi:10.1073/pnas.1615330113 (2016).

39 Akgul, G. & McBain, C. J. Diverse roles for ionotropic glutamate receptors on inhibitory interneurons in developing and adult brain. J Physiol 594, 5471–5490, doi:10.1113/JP271764 (2016).

40 He, H. Y., Hodos, W. & Quinlan, E. M. Visual deprivation reactivates rapid ocular dominance plasticity in adult visual cortex. J Neurosci 26, 2951–2955, doi:10.1523/JNEUROSCI.5554-05.2006 (2006).

41 Morales, B., Choi, S. Y. & Kirkwood, A. Dark rearing alters the development of GABAergic transmission in visual cortex. J Neurosci 22, 8084–8090 (2002).

42 Chen, J. L. et al. Clustered dynamics of inhibitory synapses and dendritic spines in the adult neocortex. Neuron 74, 361–373, doi:10.1016/j.neuron.2012.02.030 (2012).

43 van Versendaal, D. et al. Elimination of inhibitory synapses is a major component of adult ocular dominance plasticity. Neuron 74, 374–383, doi:10.1016/j.neuron.2012.03.015 (2012).

44 Fuchs, E. C. et al. Recruitment of parvalbumin-positive interneurons determines hippocampal function and associated behavior. Neuron 53, 591–604, doi:10.1016/j.neuron.2007.01.031 (2007).

45 Gu, Y. et al. Obligatory role for the immediate early gene NARP in critical period plasticity. Neuron 79, 335–346, doi:10.1016/j.neuron.2013.05.016 (2013).

46 Pelkey, K. A. et al. Pentraxins coordinate excitatory synapse maturation and circuit integration of parvalbumin interneurons. Neuron 85, 1257–1272, doi:10.1016/j.neuron.2015.02.020 (2015).

47 Liu, Y., Zhang, L. I. & Tao, H. W. Heterosynaptic scaling of developing GABAergic synapses: dependence on glutamatergic input and developmental stage. J Neurosci 27, 5301–5312, doi:27/20/5301 [pii] 10.1523/JNEUROSCI.0376-07.2007 (2007).

48 Lu, B., Wang, K. H. & Nose, A. Molecular mechanisms underlying neural circuit formation. Curr Opin Neurobiol 19, 162–167, doi:10.1016/j.conb.2009.04.004 (2009).

49 Nugent, F. S., Penick, E. C. & Kauer, J. A. Opioids block long-term potentiation of inhibitory synapses. Nature 446, 1086–1090, doi:10.1038/nature05726 (2007).

50 Gu, X., Zhou, L. & Lu, W. An NMDA Receptor-Dependent Mechanism Underlies Inhibitory Synapse Development. Cell reports 14, 471–478, doi:10.1016/j.celrep.2015.12.061 (2016).

